# OrganScaleR: An Open Shiny Tool for Principled Organ-Weight Inference in Mouse Physiology

**DOI:** 10.64898/2026.06.11.731559

**Authors:** Lucas Rebiffé, Laurent Gilquin, François Leulier, Filipe De Vadder

**Affiliations:** Institut de Génomique Fonctionnelle de Lyon, École Normale Supérieure de Lyon, CNRS UMR 5242, Université Claude Bernard Lyon 1; 69364 Lyon Cedex 07, France

**Keywords:** organ weight, allometry, ANCOVA, causal inference, Shiny application

## Abstract

**Background:** Organ weights are often divided by body weight to report “relative” organ size. Yet ratios usually stay size-dependent and become misleading. We built a simple decision path for size adjustment and wrapped it in a Shiny application so physiologists can get correct answers without coding.

**Methods:** We reused liver and body-weight data from a mouse nutrition study for confirmatory examples in two common cases: when diet groups shared a wide body-size range, and when diet produced much smaller animals with little overlap. We compared liver-to-body-weight ratios with size-adjusted regression when the body-size overlap allowed comparisons, and with causal Bayesian mediation when it did not. The full workflow was implemented in OrganScaleR, a guided R Shiny application.

**Results:** The ratio-normalized organ weight is still associated with body weight, leading to misleading comparisons. Modeling liver weight against body weight gave more cautious, size-adjusted effects when groups shared a common size range. When diets shifted body size strongly and overlap was limited, causal mediation showed that most organ differences followed the change in body weight rather than an organ-specific action. Simulations confirmed that ratios can generate false positives and biased estimates under allometric scaling, while model-based approaches remained reliable. OrganScaleR implements this decision workflow in a guided Shiny application that returns interpretable effects.

**Conclusions:** OrganScaleR selects scale, enforces common support, and routes the analysis to size-adjusted ANCOVA or causal Bayesian mediation depending on body-weight overlap. It reports adjusted effects in original units through a point-and-click workflow, removing the statistical barrier to abandoning ratio normalization.

**Highlights:**

- Body-weight ratios remain size-dependent and distort organ-weight comparisons.
- ANCOVA at a common reference body weight removes ratio bias when groups overlap.
- Causal mediation separates organ-specific from body-weight-mediated diet effects.
- Simulations confirm ratios inflate false-positive rates under allometric scaling.
- OrganScaleR guides size adjustment without coding via a point-and-click workflow.

## 1. Introduction

Organ weights are primary readouts in metabolic physiology and toxicology. They are used to infer tissue atrophy or hypertrophy, detect organ specific toxicity, and quantify the impact of dietary or pharmacological interventions. Because experimental groups often differ in body size, organ weights are still commonly expressed “per body weight” and compared as ratios. The implicit assumption is that division by body weight removes individual size differences and reveals intrinsic effects on the organ. This assumption is attractive, but it is not supported either mathematically or biologically. The idea dates to early biometric work [1–3], yet the same misconceptions persist in modern physiology [4,5].

At its core, the problem is statistical. If *Y* and *X* represent organ and body weights, their ratio *R = Y/X* is not a simple transformation with well-behaved mean and variance. There is no simple formula that expresses the mean or the variance of *R* in terms of the means and variances of *X* and *Y*, and in general *E(R)* ≠ *E (Y)/E (X)*. Even when both *X* and *Y* are independent, normally distributed variables, the distribution of *R* (the Cauchy distribution) has heavy tails and its usual moments, such as the mean and the variance, are undefined [4,6,7]. Intuitively, dividing by a variable that can be very close to zero creates occasional extremely large ratio values, which prevents the ratio from having a stable “average”. As a result, ratio statistics can behave erratically even when the underlying variables are well behaved, and familiar normal-theory intuitions about means, standard errors, and classic statistical tests become unreliable.

Biologically, ratio normalization also assumes perfect isometry, namely that organ weight scales linearly with body weight. In reality, most traits follow an allometric law,

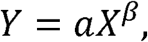

and when *β*≠ 1, the ratio *Y/X = aX*^*β*-1^ remains correlated with body weight [2,8]. This induces spurious differences between groups that merely differ in size distribution [9]. The use of ratios is therefore equivalent to fitting a misspecified model: one that omits the covariate it seeks to control [10].

Beyond scaling, ratio variables also conflate causal pathways. When a treatment affects both body weight (*B*) and an organ (*L*), the ratio *L/B* merges the direct effect of treatment on the organ with the indirect effect mediated through body weight, potentially inverting their signs [11]. The same logic explains why “percent change from baseline” performs poorly in longitudinal studies[12].

Taken together, these issues reveal that ratio normalization neither controls for size nor simplifies inference; it simply hides dependence within a distorted statistic. What is needed is an explicit model of how organ size relates to body size and how interventions act on each component.

The limitations of ratio normalization have long been recognized, and several quantitative frameworks have been developed to correct them. The central idea is that adjustment for body size or baseline values should be achieved by modelling rather than dividing. Two complementary approaches (ANCOVA or allometric regression, and causal mediation) address distinct aspects of this problem while retaining interpretability and statistical coherence.

1. *ANCOVA or allometric regression*. Rather than dividing organ weight by body weight, size adjustment should be achieved by modelling organ weight as a function of body weight and group. This approach compares groups at a shared reference body weight, while retaining the intercept and slope that describe how the organ scales with size. The resulting estimates give size-adjusted group differences in absolute organ weight and prevent the bias introduced by ratio normalization [9].
2. *Causal mediation*. When an intervention alters body weight itself, ratio methods confound direct and indirect effects. Mediation models resolve this by decomposing the total treatment effect into a direct component acting on the organ and an indirect component transmitted through body weight. Bayesian implementations offer stable inference with small samples and quantify uncertainty naturally [11].

Related approaches exist for longitudinal settings (notably Carson’s PCA-based standardization, which avoids ratio-like percent-change measures for pre/post data [5]), but fall outside the scope of the present work.

The present work addresses this methodological blind spot. We first illustrate, using simulated and empirical data, how dividing by body weight produces spurious correlations and misleading group differences. We then show how a general analytical framework combining these methods can replace ratio normalization with statistically coherent alternatives, either (i) allometric or covariance models for size correction, or (ii) causal mediation models when body weight is itself affected by treatment. Finally, we provide an open-access R Shiny application, OrganScaleR, that implements these methods in an intuitive interface, allowing physiologists to perform proper size-adjusted analyses without requiring formal statistical training. The goal is not to complicate physiology, but to replace an obsolete arithmetic shortcut with a reproducible, interpretable, and statistically sound practice.

## 2. Methods

### 2.1 Animals and diets

All animals and primary experimental procedures have been described in detail elsewhere [13]. Briefly, C57BL/6N mice were weaned at postnatal day 21 (P21) and housed with ad libitum access to food and water. From P21 to P56, mice were fed either a control diet (CD) or an isocaloric low-protein diet (LPD) differing only in protein content. At P56, mice were fasted for 6 hours in the morning, anaesthetized and euthanized for organ collection. Body weight and liver weight were recorded on the same day. Body composition (including lean mass) was assessed non-invasively by low-field NMR at P38, prior to tissue collection [13]. For the present methodological study, we used liver and body weight data from two experimental batches of females and two batches of males. These datasets were chosen as illustrative examples of common situations in physiology: (i) moderate differences in body weight with broad overlap between groups (females), and (ii) large diet-induced differences in body weight with limited overlap (males). Here, males and females were analyzed separately. A third dataset, comprising both sexes in a balanced 2×2 design (Section 2.2), was used solely to illustrate the application’s behavior in factorial settings. The goal was to illustrate analytical issues with ratio normalization and their remedies; the factorial dataset demonstrates stratified analysis rather than a formal test of Sex×Diet interaction.

### 2.2 Empirical datasets

For females, we analyzed liver and body weight from adult mice (P56) fed CD or LPD for 5 weeks (n = 18 CD and 21 LPD). This dataset had broadly overlapping body-weight distributions between diets and was used to illustrate classical ANCOVA and allometric regression as alternatives to liver-to-body-weight ratios.

For males, we used an analogous dataset from adult CD- and LPD-fed mice (n = 18–19 per diet) in which the LPD produced a marked reduction in body weight and thus limited overlap between groups. This dataset was used to illustrate Bayesian causal mediation when body weight is both an outcome of diet and a mediator of liver weight.

For the 2×2 example, we used lean mass and body weight from a balanced factorial subset of the same cohort, comprising both sexes and both diets (n = 10 per Sex × Diet group, 40 mice total). Because the four groups occupied largely non-overlapping body-weight ranges, no single body weight was shared across all groups; this dataset was therefore used to illustrate stratified causal mediation, in which the diet effect on lean mass is decomposed within each sex.

### 2.3 Ratio variables and simple group comparisons

In all datasets, liver-to-body-weight ratios were computed as:

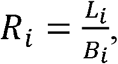

where *L*_*i*_ is liver weight and *B*_*i*_ is body weight for mouse *i*. For descriptive purposes, we summarized body weight, liver weight and the ratio by diet (mean ± SD) and inspected their distributions. In the female and male examples, normality within diet and equality of variances for the ratio were assessed with Shapiro–Wilk tests and F-tests, respectively. We then performed two-sample *t*-tests on the ratio *R* to mimic common practice and to contrast these results with model-based analyses.

To evaluate whether the ratio truly removed dependence on body size, we fitted a linear model of the form

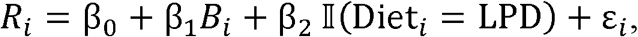

where *ε*_*i*_ are residual errors and 𝕀(·) is an indicator variable. A significant slope *β*_1_ indicates that the ratio remains size-dependent.

### 2.4 Allometric and ANCOVA models (female example)

For the female dataset, where CD and LPD body-weight distributions overlapped, we analyzed liver weight directly as a function of body weight and diet. We fitted two competing size-adjustment models. The linear ANCOVA on the original scale was

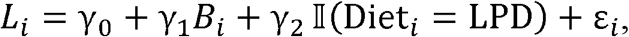

and the log–log allometric model was

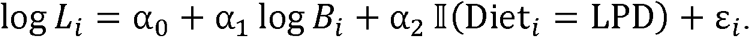

Here *L*_*i*_ is liver weight, *B*_*i*_ is body weight, and 𝕀(·) codes the LPD group. In both models we inspected standard residual diagnostics (residuals versus fitted values, Q–Q plots) to verify approximate normality and homoscedasticity.

Model choice was based on predictive performance rather than goodness-of-fit alone. We compared models by AIC and by cross-validated RMSE computed on the original liver-weight scale. Specifically, we performed stratified K-fold cross-validation (K = 5, folds balanced by diet), refitting each model on the training folds and predicting the held-out fold. For the log–log model, predictions were back-transformed using Duan’s smearing estimator calculated on training residuals to reduce re-transformation bias before computing RMSE. The scale (raw vs log-log) with the lower cross-validated RMSE was selected for primary reporting, while the alternative parametrization was retained as a sensitivity check.

To prevent adjusted comparisons outside the region of shared body-weight support, all size-adjusted effects were evaluated at a reference body weight *B*^*^ defined within the trimmed overlap of the two diets. We first estimated, for each diet, the 5th and 95th percentiles of body weight, then defined the overlap window as the intersection of these percentile ranges. If the overlap window was empty, adjusted ANCOVA contrasts were considered invalid and inference would be routed to causal mediation. When overlap existed, *B* was set to the geometric mean body weight of individuals lying within the overlap window.

Size-adjusted marginal means 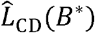 and 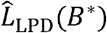 and their 95% confidence intervals were obtained using the R package emmeans. To mitigate potential mild heteroscedasticity, contrasts and intervals were computed with HC3 robust covariance estimators. For log–log models, estimated marginal means and intervals were re-expressed on the original liver-weight scale after smearing-based back-transformation. These adjusted means were displayed on top of the raw liver-weight distributions to show both the observed data and the size-standardized estimates.

### 2.5 Simulation study of ratio versus model-based methods

To explore the behavior of organ-to-body-weight ratios under controlled conditions, we simulated datasets from a simple allometric relationship calibrated on the empirical female liver–body data.

Body weight *B* was generated from group-specific normal distributions, and liver weight *L* was generated from a log–log allometric model of the form

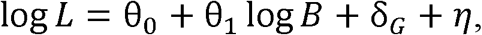

where *θ*_0_ and *θ*_1_ define the baseline allometric relationship, *δ*_*G*_ is a group effect on liver weight, and η, is a normally distributed residual term. Parameters were chosen so that the body-weight range and variability matched the empirical data, and the allometric exponent *θ*_1_ was approximately 1.6. We considered three scenarios (20 animals per group, 5,000 replicates per scenario):

1. Null scenario: groups differ only in body-weight distribution; *δ*_*G*_ *=0* and the true difference in liver weight at equal body weight is zero.
2. Direct-effect scenario: groups have identical body-weight distributions, but a positive organ-specific effect is imposed via *δ*_*G*_, yielding a true between-group difference of about 0.18–0.20 g at equal body weight.
3. Combined scenario: both a positive direct effect on liver weight and a shift in body-weight distribution are present.

For each simulated dataset, we estimated the between-group difference in liver weight using:

- a t-test on the ratio *R = L/B*,
- a linear ANCOVA of the form *L*_*i*_ *= y*_0_ *+ y*_1_ *B*_*i*_ *+ y*_2_*G*_*i*_ *+ E*_*i*_
- a log–log allometric model of the form *logL*_*i*_ *= a*_0_ *+ a*_1_*logB*_i_ *+ a*_2_*G*_*i*_ *+ E*_*i*_, where *G*_*i*_ denotes group.

For each method in each scenario, we summarized across simulations (i) the mean estimated group difference at equal body weight (bias), (ii) the root mean square error (RMSE) of this estimate, and (iii) the empirical rejection rate of a null hypothesis of no difference at a 5 % significance level (type I error in the null scenario, power in the non-null scenarios).

### 2.6 Bayesian mediation analysis (male example)

In the male dataset, the low-protein diet produced a large reduction in body weight and only a narrow region of overlap between CD and LPD body-weight distributions. Although a trimmed overlap was still detectable, it was confined to a small interval, so any ANCOVA or log–log comparison at a single reference body weight would be driven by a minority of animals and would answer a question that is not representative of either group. Following Lazic et al. [11], we therefore treated body size as a mediator of the diet effect on liver weight and used a causal mediation model as the primary analysis.

Let *D*_*i*_ denote diet (CD or LPD), *B*_*i*_ body weight, and *L*_*i*_ liver weight for mouse *i*. To reduce artefactual coupling between mediator and outcome when the organ contributes non-negligibly to body mass, the mediator was defined as organ-free body weight

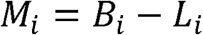

We assumed a simple mediation structure in which diet affects organ-free body weight, organ-free body weight affects liver weight, and diet may also exert a residual direct effect on liver weight after accounting for body size. This was implemented as a Bayesian multivariate model with the two following Gaussian link functions:

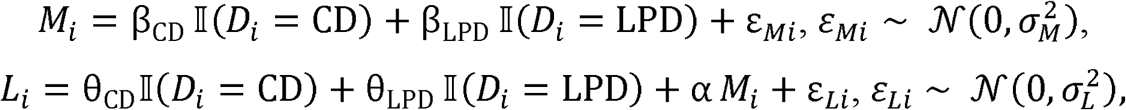

with independent normal residuals *ε*_*Mi*_ and *ε*_*Li*_. The organ weights and body weights are each assumed to have equal variance for the two diets. The variances are respectively modeled by 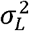 and 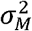. This decomposition assumes sequential ignorability: conditional on diet, there are no unmeasured confounders of the organ-free body weight–liver weight relationship. This assumption is supported here by the randomized, single-strain design with uniform housing and age at sacrifice, which limits sources of uncontrolled variation between the mediator and outcome.

From the joint posterior of the regression coefficients, we decomposed the diet effect on liver weight (LPD minus CD) into:

- Direct effect (DE): *DE = θ*_*LPD*_ *- θ*_*CD*_, representing the part of the liver difference not explained by body weight;
- Indirect effect (IE): *IE = a(β*_*LPD*_ *- β*_*CD*_), representing the component transmitted through diet-induced changes in body weight;
- Total effect (TE): *TE = DE + IE*, corresponding to the overall difference in liver weight between LPD and CD males.

The model was fitted using the R package brms (which uses Stan as a backend) with weakly informative priors on regression coefficients and residual standard deviations, four MCMC chains, 4,000 iterations per chain (2,000 warm-up), and NUTS (No-U-Turn-Sampler) tuning parameters set to improve stability `(adapt_delta = 0.99, max_treedepth = 15)`. Convergence was assessed using 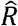 and effective sample sizes, and posterior predictive checks were inspected for both mediator and outcome models.

For each effect (TE, IE, DE), we reported the posterior mean, 95% credible interval, the posterior probability of a negative effect (LPD < CD), and the probability of lying within a Region Of Practical Equivalence (ROPE). The ROPE was defined as ±5% of mean liver weight in CD males, so effects inside this interval were considered physiologically negligible. To verify that conclusions did not depend on any arbitrary reference weight, we performed standardization with g-computation on the fitted mediation model. For a subset of posterior draws, we simulated organ-free body weights from the CD mediator distribution,

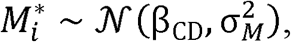

then simulated liver weights for both diets at these 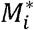,

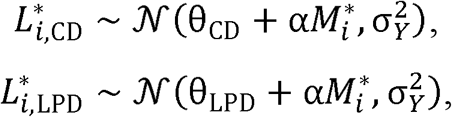

and computed the mean difference 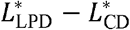 for each posterior draw. This quantity represents the expected diet effect on liver weight if both groups shared the control distribution of organ-free body weight. Its posterior mean, 95% credible interval, and ROPE probability were compared with the coefficient-based direct effect as a consistency check.

### 2.7 OrganScaleR Shiny application for principled organ-weight analysis

To facilitate routine use of the model-based alternatives to ratio normalization developed here, we developed an open-access R Shiny application that implements the analytical workflow in an interactive interface. The application is intended to reproduce the present framework without adding new statistical procedures.

#### 2.7.1 Data input and experimental design

Users upload a CSV or Excel table containing organ weight, body weight, and one or two categorical grouping variables. Organ and body-weight units are specified by the user and are converted internally to grams for modeling. All reported estimates, confidence intervals, and plots are returned in the original units. When a single factor is provided, it is analyzed as a one-factor design. When two factors are provided, the Shiny application constructs a composite group corresponding to the full 2 × 2 design (Factor 1 × Factor 2) and stores both factor labels for downstream contrasts.

#### 2.7.2 Automatic choice between RAW ANCOVA and log–log allometry

The application fits both (i) a linear ANCOVA model on the original organ-weight scale and (ii) a log–log allometric model, each with organ weight as the response, body weight as the continuous explanatory variable, and group(s) as categorical explanatory variables. Scale selection is based on stratified K-fold cross-validated RMSE (default K = 5, folds balanced across group levels), computed on the original organ-weight scale. For log–log models, predictions and adjusted means are back-transformed using Duan’s smearing estimator, computed either globally or by group, to avoid retransformation bias. Users may override the automatic choice in Expert mode, but the cross-validated metric is always displayed for transparency.

#### 2.7.3 Guardrails against extrapolation and definition of the reference body weight

To prevent adjusted comparisons outside the support of the data, the Shiny application detects the shared body-weight range across groups using a trimmed overlap rule defined as the intersection of the 5th to 95th percentile ranges in each group. Adjusted group means and contrasts are then computed at a reference body weight defined as the geometric mean body weight of animals lying within this overlap window. When no trimmed overlap exists, global ANCOVA contrasts are disabled and the user is directed to causal mediation (below), reflecting the framework of the manuscript.

#### 2.7.4 Factorial designs and simple effects

In 2 × 2 designs, the Shiny application first fits the full model with a slope-by-group interaction. The interaction is retained only if its F-test *P*-value is below 0.01; otherwise, parallel slopes are assumed and the reduced model is used for inference. Adjusted means are estimated with the R package emmeans at the overlap-based reference body weight using HC3 robust covariance estimators to mitigate mild heteroscedasticity. Post-hoc comparisons can be performed as Dunnett-type tests against a control level (based on what the user chose when loading the data), or as fully pairwise tests using Tukey or Holm adjustment. For users requesting simple effects in 2 × 2 designs, contrasts within each level of the other factor are computed at the same reference body weight, preserving interpretability on a common size scale.

#### 2.7.5 Pairwise and “soft” matching options

By default, contrasts use the global overlap window shared across all groups. Users can optionally request pairwise overlap, in which each contrast is evaluated at its own overlap-based reference body weight. The Shiny application also exposes a “soft” matching option that allows a small user-defined body-weight gap (default 5%) to be bridged by evaluating at the nearest common body weight. This mode is explicitly flagged in outputs because it weakens the common-support guarantee.

### 2.7.6 Causal ANCOVA (Expert mode)

When body weight is affected by treatment, the application provides an Expert-mode Bayesian causal mediation module matching the model specified in the manuscript. Within the interface this module is labelled “Causal ANCOVA”. The mediator can be set to raw body weight or to “organ-free body weight” (body weight minus organ weight), the latter being the default to limit artefactual coupling when the organ constitutes a non-negligible fraction of body mass. In one-factor settings, mediation is only enabled when Factor 1 has exactly two levels. In 2 × 2 settings, mediation is computed for Factor 1 within each level of Factor 2, provided all four cells are present. The multivariate Gaussian system is fit with the R package brms with weakly informative priors, four MCMC chains (4,000 iterations each, 2,000 warm-up), and standard convergence and posterior predictive checks. Posterior draws yield total, direct, and indirect effects, each summarized by posterior mean, 95% credible interval, posterior probabilities of being negative or positive, and Region of Practical Equivalence (ROPE) probability. The ROPE is defined as ±5% of the mean control organ weight, computed within the relevant baseline stratum (overall for one-factor mediation, or within each Factor-2 level in 2 × 2 mediation). To evaluate sensitivity to reference body-weight choices, the Shiny application performs a g-computation check by simulating organ weights for both groups under the control body-weight distribution, yielding a standardized direct effect with the same posterior summaries and ROPE probability.

### 2.7.7 Plain-English summaries

To facilitate interpretation by non-specialists, the application generates a structured plain-English narrative alongside each result panel. These summaries do not perform additional analyses. They restate the numerical outputs already reported in the tables and plots. In ANCOVA or allometry mode, the narrative reports the overlap-based reference body weight, per-group adjusted means at that reference in original units, the corresponding contrasts with confidence intervals, overall model fit (R^2^), effect sizes (η^2^) for the covariate, grouping factor, and interaction, and a qualitative descriptor of the strength of evidence based on the *P*-values. In causal ANCOVA mode, the narrative restates the causal question and comparison, specifies the mediator choice and ROPE scale, and summarizes total, indirect, and direct effects with credible intervals and their interpretation relative to the ROPE, with an optional g-computation sensitivity statement when available.

### 2.7.8 Diagnostics and error handling

All steps write structured INFO, WARN, and ERROR messages to a Diagnostics tab and to the console. A global Shiny error hook captures uncaught failures and surfaces them to the user, preventing silent return of results.

### 2.8 Software and reproducibility

All analyses and the Shiny application were developed in R v4.4.3. Model fitting relied primarily on the R package brms (v2.23.0) for Bayesian mediation, `emmeans` (v1.11.2-8) for adjusted means and contrasts, and `sandwich` (v3.1-1) for robust covariance estimation. Data handling and plotting used the R packages `dplyr` (v1.1.4) and `ggplot2` (v3.5.2). A complete package list and session information are provided in the code repository.

## 3. Results

### 3.1 Organ-to-body weight ratios look well-behaved but remain strongly body weight-dependent

Because organ weights are often expressed relative to body weight, we first examined whether this conventional ratio-based normalization was appropriate in a simple situation where body weight was similar between groups. For this illustrative analysis, we used publicly available data (available on Zenodo, see Data availability) from adult female mice (56 days old) drawn from two experimental batches and fed for 5 weeks with either a control diet (CD) or a low-protein diet (LPD), under the same experimental conditions as described previously [13]. This dataset was selected because body weight distributions were broadly overlapping between CD and LPD females (**Figure 1A**).

**Figure 1.**
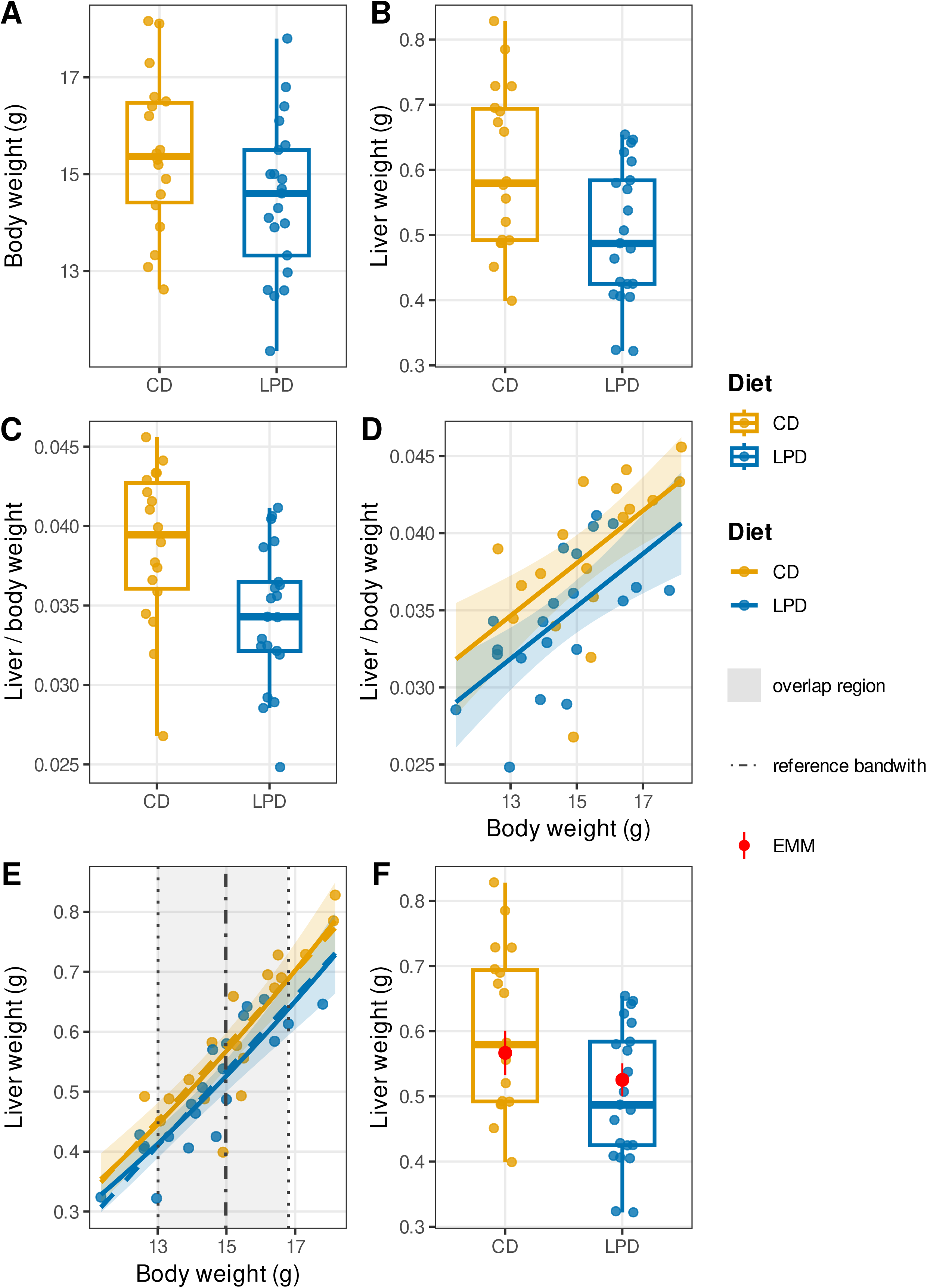
Ratio-based “normalization” does not remove body-weight scaling of liver weight, motivating covariate-adjusted comparisons. **(A–C)** Boxplots with individual mice for body weight **(A)**, liver weight **(B)**, and liver-to-body-weight ratio **(C)** in CD vs LPD females. The unadjusted ratio differs between diets **(C)**, two-sample *t*-test *P* = 0.005. **(D)** Linear regression of liver-to-body-weight ratio vs body weight shows the ratio remains strongly dependent on body weight (Body weight term *P* = 8×10^-5^). **(E)** Linear ANCOVA on the raw scale (dashed line) and log-log allometry (solid line) give similar qualitative conclusions. Vertical lines indicate the overlap-based body-weight window and the reference body weight used for adjusted means. **(F)** Liver-weight boxplot with covariate-adjusted estimated marginal means at the overlap-based reference body weight (red dots with 95% CI); the adjusted CD–LPD contrast is weaker than the ratio *t*-test (*P* = 0.052).

Mean body weight was slightly higher in CD than in LPD females, but with comparable variability (15.4 ± 1.6 g vs. 14.5 ± 1.6 g). Liver weight was lower in LPD females (0.50 ± 0.11 g) than in controls (0.60 ± 0.13 g), and the corresponding liver-to-body-weight ratio was also reduced (0.034 ± 0.004 vs 0.039 ± 0.005) (**Figure 1A-C**). Within each diet, body weight, liver weight and their ratio were approximately normally distributed, and the variance of the ratio did not differ between groups (all Shapiro–Wilk *P* ≥ 0.05; F-test on the ratio *P =* 0.61). Under these conditions, a two-sample *t*-test on the ratio would usually be considered acceptable and indeed detected a significant difference between diets (Δ = 0.0044 in liver/body ratio; *P =* 0.005, **Figure 1C**). Taken at face value, this analysis would suggest that females on the control diet have a larger “relative liver size” than those on the low-protein diet.

To test whether this ratio removed the dependence on body weight, we next regressed the ratio on body weight and diet (**Figure 1D**). In this model, the ratio increased with body weight within diets and still differed between diets. In other words, heavier mice within a given diet have a larger liver-to-body-weight ratio, so the ratio is not independent of body weight, the covariate it is supposed to remove. Body weight remained a strong predictor of the liver-to-body-weight ratio (partial η^2^ = 0.35, *P* = 8.3×10^-5^), indicating that ratio normalization failed to remove size dependence. The significant *t*-test on ratios therefore reflects both the diet effect and residual scaling with body weight.

We therefore modelled liver weight directly as a function of body weight and diet. In a linear ANCOVA, liver weight increased with body weight and remained lower in LPD females after adjustment for body weight. A log–log allometric model, where both liver and body weight were analyzed on a logarithmic scale, led to the same qualitative conclusion (**Figure 1E**). Over this restricted size range, the linear model was favored by AIC (−108 vs. −56) and cross-validated RMSE was marginally lower for the log–log model (0.0578 vs. 0.0582 g). Because cross-validated prediction errors were nearly identical, we retained the linear ANCOVA as the primary model for its simpler interpretation on the original scale, and report the log–log results as a sensitivity check.

To express the diet effect in interpretable units without extrapolation, we estimated liver weight for both diets at a common reference body weight defined within the trimmed overlap of body-weight distributions (**Figure 1F**). The overlap window was 13.0 to 16.8 g, yielding a reference body weight of 15.0 g (geometric mean within overlap). At this reference, adjusted liver weight was 0.575 g in controls (95% CI 0.543 to 0.607 g) and 0.533 g in LPD females (95% CI 0.508 to 0.558 g) under linear ANCOVA, corresponding to an adjusted difference of −0.042 g. With robust HC3 uncertainty, this contrast was borderline (P = 0.052). The log–log model gave nearly identical adjusted means (0.567 vs. 0.525 g) and the same effect size (−0.041 g), with slightly wider robust uncertainty (P = 0.065). These adjusted means and their 95% confidence intervals are plotted on top of the raw box plots of liver weight (**Figure 1F**). Overall, even when ratio values are approximately normal and have similar variance, a *t*-test on organ-to-body-weight ratios can be misleading because the ratio often remains correlated with body weight. Modelling the organ directly as a function of body weight and treatment and comparing groups at a shared reference body weight within common support, yields an effect estimate with transparent dependence on size and avoids the false certainty induced by ratio normalization. In these females, the adjusted diet effect remains close to −0.04 g, but the robust, overlap-restricted analysis makes clear that the evidence is weaker than suggested by the ratio *t*-test alone.

### 3.2 Organ-to-body weight ratios fail under allometric scaling

To assess how organ-to-body weight ratios behave under controlled conditions, we simulated datasets from a simple log–log allometric model calibrated on the liver–body relationship of our empirical example (exponent 1.6, n = 20 per group). We considered three scenarios: (i) a null scenario where groups differed only in body weight, (ii) a scenario with a positive direct effect on organ weight but identical body weight distributions, and (iii) a scenario combining a positive direct effect with a shift in body weight. Each scenario was replicated 5,000 times, and for each dataset we estimated the between-group difference in organ weight using the ratio, a size-adjusted ANCOVA, and a log–log allometric model.

Figure 2 summarizes what would happen if the same experiment were repeated many times. For each scenario and each method, the violin plots show the distribution of 5,000 estimated between-group differences in organ weight. The horizontal red line marks the *true* difference in organ weight between groups at the same body weight, as specified by the simulation model. **Table 1** reports the corresponding numerical summaries.

In the null scenario, the two groups were generated to have identical organ size at any given body weight and differed only in body weight. In that case, the true difference at equal body weight is zero (red line at 0). Nevertheless, the ratio method declared a “significant” group difference in 36 % of the simulated experiments (type I error 0.36; **Table 1**), even though its estimates cluster visually around zero. In contrast, the ANCOVA and log–log allometric approaches, which explicitly estimate organ size conditional on body weight, rejected the null in about 5 % of simulations, as expected for a 5 % significance level.

**Figure 2.**
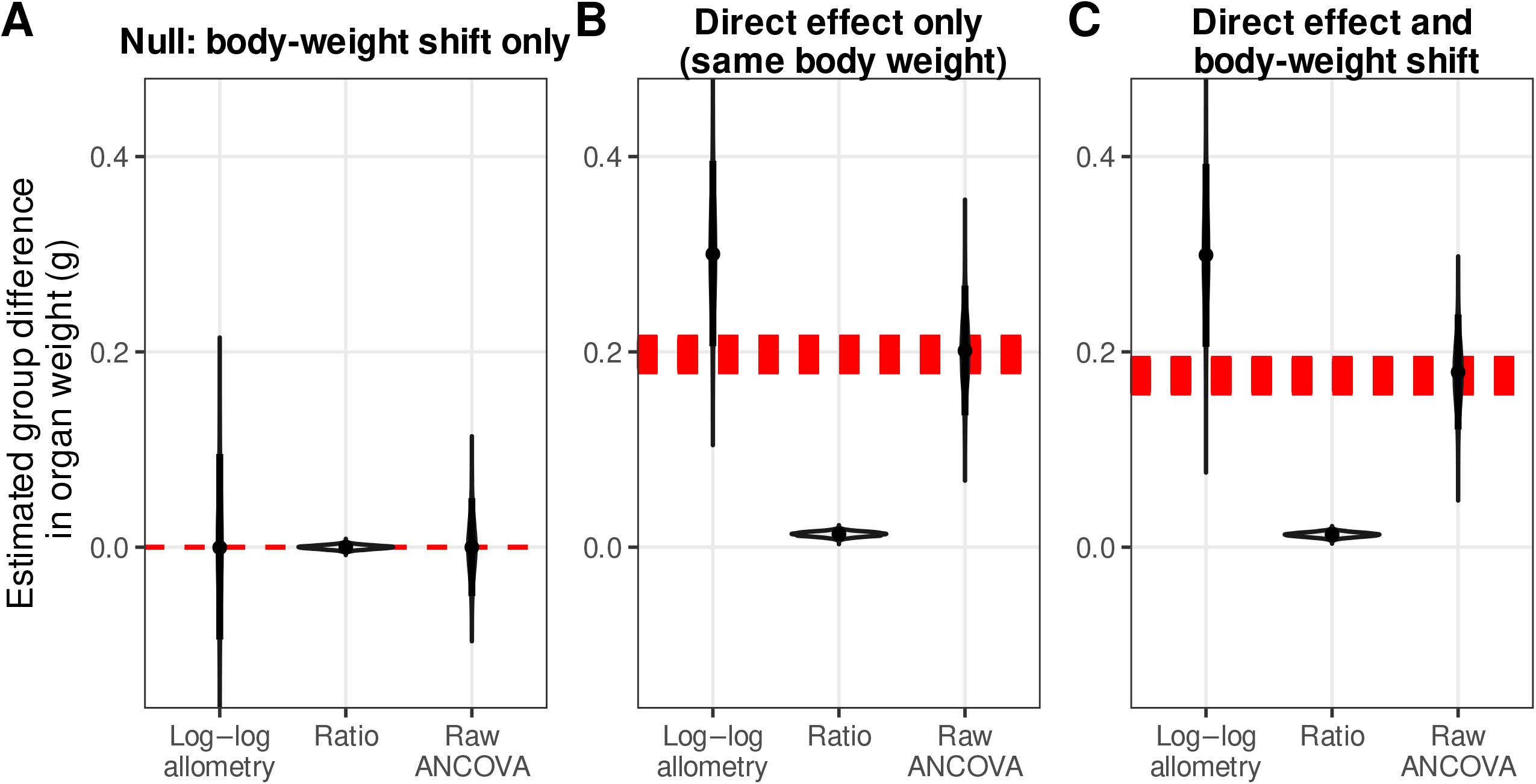
Simulations show organ-to-body-weight ratios fail under allometric scaling. **(A-C)** Results of 5,000 simulations of allometric organ scaling under three scenarios: **(A)** null organ effect with a body weight shift only; **(B)** direct organ effect only (no body-weight shift), where groups differ in organ weight at the same body size; **(C)** direct organ effect plus a body-weight shift. Violin plots show the distribution (across simulated datasets) of the estimated group difference in organ weight (group B − group A) obtained with organ-to-body-weight ratio, raw ANCOVA, or log–log allometry. Points indicate the median estimate and vertical bars the central quantile interval (as defined in the plot). The dashed horizontal line indicates the true effect used to generate the data. All panels use the same y-axis scale. Detailed simulation results are provided in **Table 1**.

In the two non-null scenarios, we deliberately set the model so that, for animals of the same body weight, organs in group B were about 0.18–0.20 g heavier than in group A (red line near 0.2 g, **Figure 2B-C**). The ratio method almost completely missed this effect. On average, its estimates were 0.17–0.18 g below the true value, that is, it “saw” a difference close to 0 g instead of 0.2 g. The typical size of the error (RMSE) was also 0.17–0.18 g, meaning that in a representative experiment the ratio-based estimate was wrong by about 0.17 g, which is almost as large as the true effect itself (**Table 1**). As a result, in **Figure 2B-C** the black shapes for the ratio are centered near 0 while the red line marking the true effect sits around 0.2 g. By contrast, the ANCOVA and log–log allometric models recovered the intended difference at equal body weight. Their average error was essentially zero (about 0.003–0.004 g, **Table 1**), and their typical error was much smaller (RMSE 0.03–0.04 g, roughly five to six times smaller than for the ratio). In other words, these model-based approaches not only point to the correct effect size but also do so with much less scatter across repeated experiments, yielding power above 99 %. Of the two model-based approaches, raw ANCOVA was essentially unbiased (mean error < 0.005 g), whereas the log–log model showed moderate residual bias (0.10–0.12 g), consistent with retransformation error when the allometric exponent departs substantially from unity.

### 3.3 Bayesian mediation allows comparison of organ weights when overlap is narrow and treatment shifts body weight

The previous examples assumed reasonably similar body-weight distributions between groups, so that liver weights could be compared at a common reference body weight using ANCOVA or allometric models. We next considered an independent illustrative dataset from adult male mice, chosen because the low-protein diet produced a strong reduction in body weight and therefore only a narrow region of overlap between groups (**Figure 3A,D**). Adult males fed CD weighed 20.0 ± 2.5 g on average, whereas males on LPD weighed 14.9 ± 2.3 g (mean ± SD, n = 19 and 18, respectively). Liver weight was also lower in LPD males (0.510 ± 0.176 g) than in controls (0.864 ± 0.257 g), and the conventional liver-to-body-weight ratio was reduced from 0.043 ± 0.009 in CD to 0.033 ± 0.008 in LPD animals (**Figure 3B-C**).

**Figure 3.**
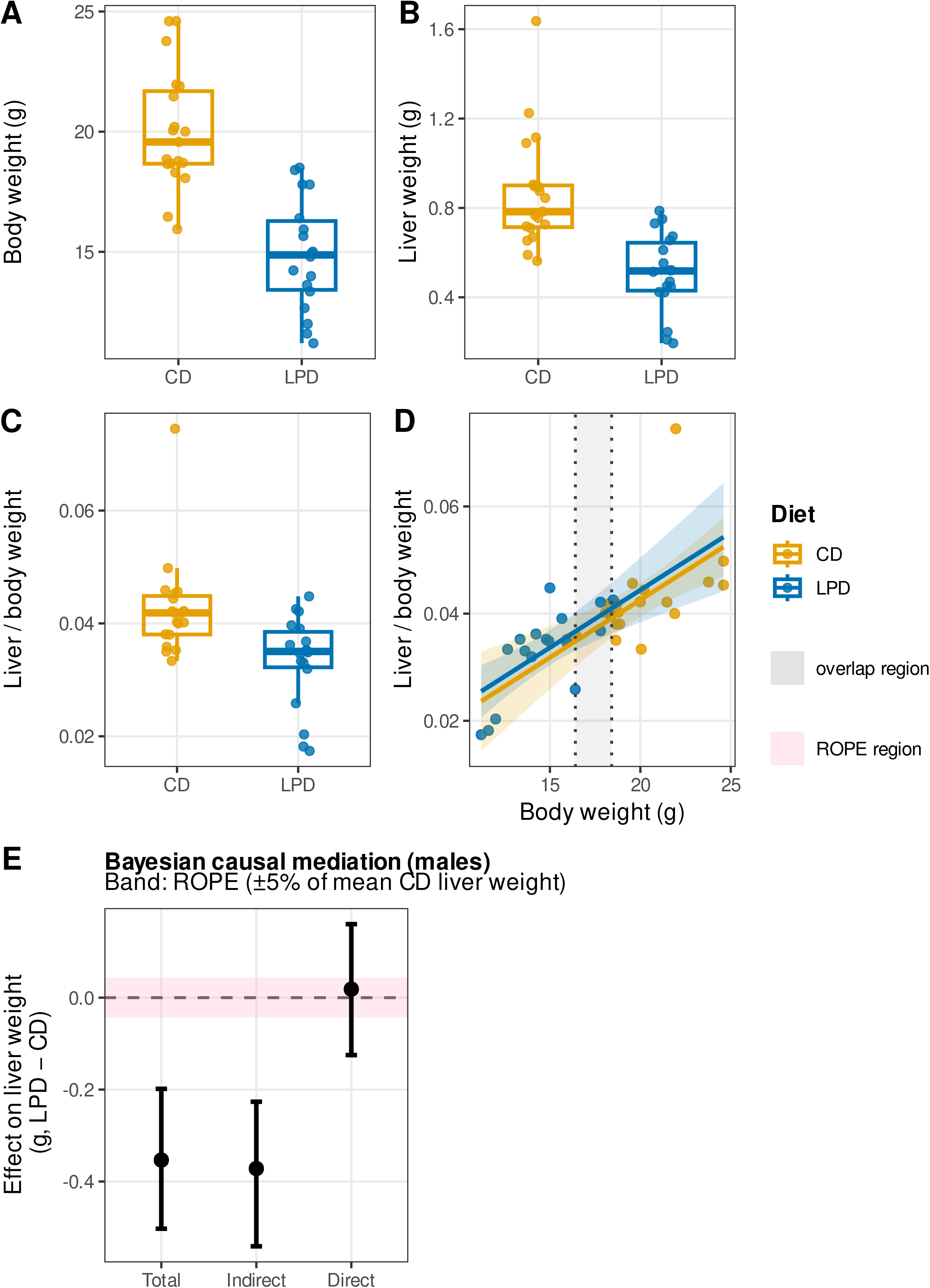
When body-weight overlap is limited, Bayesian causal mediation separates body-size effects from organ-specific effects of diet. **(A–C)** Boxplots with individual mice for body weight **(A)**, liver weight **(B)**, and liver-to-body-weight ratio **(C)** in CD vs LPD males. The unadjusted ratio differs between diets **(C)**, two-sample *t*-test *P* = 0.002. **(D)** Linear regression of liver-to-body-weight ratio vs body weight shows a strong body-weight dependence (Body weight term P = 9×10^-5^). Vertical lines indicate the trimmed body weight overlap window (5^th^–95^th^ percentile intersection), highlighting the limited common support between diets. **(E)** Mediation schematic: diet shifts organ-free body weight (body size excluding liver), which scales liver weight; a separate path represents any diet effect on liver weight beyond body size (organ-specific). **(F)** Bayesian causal mediation decomposition of the LPD−CD difference in liver weight into Total, Indirect (via body weight), and Direct (organ-specific at the same body weight) effects (dots with 95% credible intervals). Posterior probabilities for each effect being negative are reported, and the negligible-effect band (ROPE) is highlighted in grey.

Despite superficially “clean” summary statistics, several assumptions behind ratio testing fail here. In CD males, liver weight and the ratio deviate from normality (Shapiro–Wilk P = 0.008 and 8.4 × 10^-5^, respectively), although the ratio variances remain comparable between diets (F-test *P* = 0.72). A two-sample *t-*test on the ratio would still be routinely applied and yields a significant difference (Δ = 0.0092 in liver/body ratio, *P* = 0.002), but this *P*-value is not trustworthy as evidence for a liver-specific diet effect given the non-normal ratios and, more importantly, the strong body-size shift induced by diet (**Figure 3C**).

As in females, we asked whether the liver-to-body-weight ratio really removed the effect of body size. Regressing the ratio on body weight and diet (`ratio ~ Body_weight + Diet`) showed that the ratio increases with body weight within diets, while the additional diet term contributes little once body weight is included. Thus, “normalizing” by body weight does not eliminate size dependence, and the ratio difference largely reflects the fact that LPD males are lighter, not an independent liver effect (**Figure 3D**).

The trimmed overlap of body weights between diets was small (16.4 to 18.4 g). Any ANCOVA or log–log comparison at a single reference weight would therefore answer a narrow, potentially unrepresentative question. To deal with this situation, we implemented Bayesian causal mediation [11] as a multivariate Gaussian model with two linked equations. In this mediation analysis, we treated organ-free body weight as the mediator, defined as total body weight minus liver weight for each mouse. Using organ-free rather than total body weight avoids the trivial situation where the liver partly “predicts itself” through its own contribution to body weight, which can create artificial coupling when the organ represents a non-negligible fraction of total mass.

We then fitted two linked regression models. The first model described how organ-free body weight depends on diet (CD or LPD). The second model described how liver weight depends jointly on diet and organ-free body weight. This set-up corresponds to a simple causal diagram where diet changes body size, body size influences liver size, and diet may also have an additional direct effect on liver beyond its effect on body size (**Figure 3E**).

Using Bayesian inference, we obtained for each quantity a distribution of plausible values. From these we decomposed the effect of LPD versus CD on liver weight into three components, all expressed in grams: (i) the total effect (TE), which is the overall difference in liver weight between the two diets; (ii) the indirect effect (IE), which is the part of this difference that operates through changes in body size; and (iii) the direct effect (DE), which is the remaining difference in liver weight after taking body size into account. For each effect we report the posterior mean, the 95% credible interval (range containing the 95% most plausible values), and the probability that the effect lies within a Region Of Practical Equivalence (ROPE). Here the ROPE was defined as ±5% of mean liver weight in CD males (±0.043 g), so effects inside this interval are considered physiologically negligible.

The results showed a large overall loss of liver weight in LPD males, with a total effect of approximately −0.37 g relative to CD. Almost all this loss was explained by body size: the indirect effect through organ-free body weight was about −0.38 g, clearly outside the ROPE, whereas the direct effect was very small, close to +0.01 g on average and largely contained within the ROPE. In practical terms, once the strong reduction in body size is accounted for, there is no convincing evidence for an additional liver-specific action of the low-protein diet in males (**Figure 3F**).

The mediation approach provides a clear answer in a situation where classical size adjustment becomes fragile. When body weight is strongly shifted and overlap is narrow, comparing groups at a single “common” weight is either dominated by a handful of animals or forces extrapolation. Mediation avoids that trap by using the whole dataset, separating what is explained by body size from what is not, and quantifying the uncertainty on each component. In physiology, this means being able to state plainly whether an organ change is mostly a consequence of smaller animals or whether there is credible evidence for an organ-specific effect, without relying on ratios that remain size-dependent and can exaggerate significance.

### 3.4 OrganScaleR Shiny application eases the decision framework and prevents ratio-style extrapolation

To facilitate routine application, we implemented the workflow described above in an open-access R Shiny interface, called OrganScaleR (**Figure 4A**). We illustrate its behavior on the same female and male liver datasets, and on an additional 2 × 2 lean mass dataset, emphasizing the estimands produced by each branch of the decision tree.

**Figure 4.**
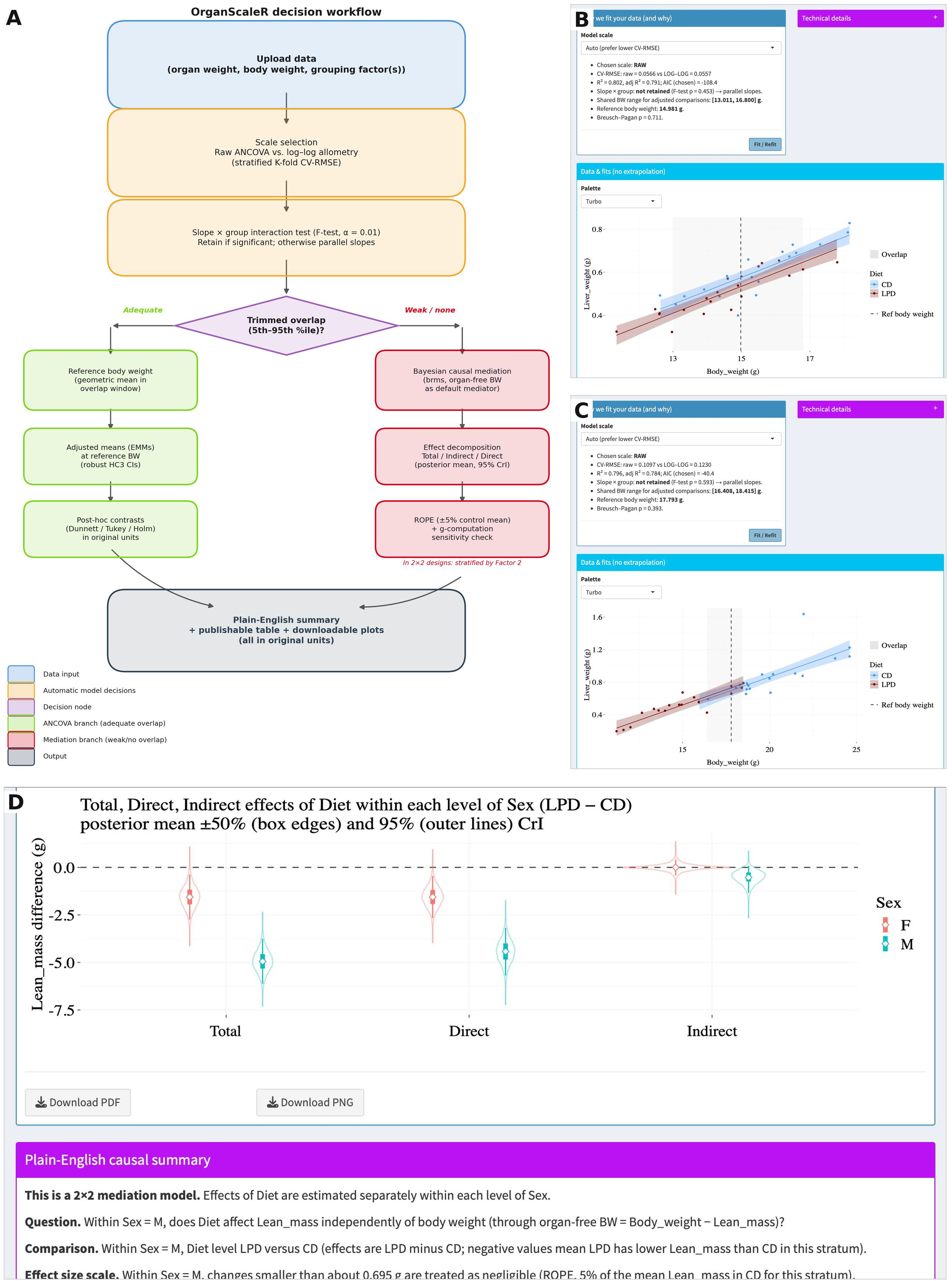
OrganScaleR implements a guided decision workflow for principled organ-weight analysis. **(A)** Decision flowchart. Users upload organ weight, body weight, and grouping factor(s). The application selects between raw ANCOVA and log–log allometry by stratified cross-validated RMSE, tests whether slopes differ across groups, and detects shared body-weight support via a trimmed overlap rule (5th–95th percentile intersection). When overlap is adequate (left branch), adjusted means are estimated at a data-derived reference body weight with robust confidence intervals and post-hoc contrasts in original units. When overlap is weak or absent (right branch), Bayesian causal mediation decomposes the treatment effect into total, indirect (via body weight), and direct (organ-specific) components, evaluated against a Region Of Practical Equivalence (ROPE). In 2×2 factorial designs, mediation effects are estimated within each level of the second factor. Both branches converge on a plain-English summary with downloadable plots and tables. **(B)** Female liver dataset, model-fitting step. The broad body-weight overlap defines a valid reference body weight (dashed line), routing the analysis to the ANCOVA branch; the plot shows group-specific fits and the shaded overlap region. **(C)** Male liver dataset, model-fitting step. The control and low-protein groups occupy largely distinct body-weight ranges, leaving only a narrow overlap; adjusted comparisons would require extrapolation, routing the analysis to the causal-mediation branch. **(D)** 2×2 factorial dataset (Diet × Sex), stratified causal mediation. With no body-weight range shared across all four groups, global ANCOVA is disabled and the diet effect (LPD − CD) is decomposed into total, indirect, and direct components separately within each Sex stratum (F, M). Points are posterior means; box edges and outer lines denote 50% and 95% credible intervals. A substantial direct effect persists in both sexes, indicating an organ-specific effect of diet on lean mass beyond body-weight differences.

In the female dataset, users upload the table, select the organ and body-weight columns, and define Diet as the single grouping factor, with CD as control (**Figure 4B**). The application then fits both the raw ANCOVA and log–log allometric models and displays their stratified cross-validated RMSE on the original organ-weight scale. For this dataset, prediction errors were nearly identical across scales, and the application selected the raw model by default, while making the scale comparison explicit (**Figure 4B**). Consistent with the framework of the manuscript, it detected a trimmed shared body-weight window across diets and computed an overlap-based reference body weight within it. Adjusted liver means at this reference were returned in original units together with robust 95% confidence intervals and a Diet contrast evaluated at the same reference size (**Figure 4B**). These adjusted means were plotted on top of the raw data, and the corresponding numerical results were provided as a ready-to-report output table (**Figure 4B**). Thus, for a one-factor design with adequate overlap, the application reproduces the main ANCOVA/allometry branch while enforcing common support and reporting effects directly in grams rather than ratios.

In the male dataset, where Diet produced a strong shift in body weight, the application again performed transparent scale comparison and selected the RAW scale but detected that the trimmed overlap of body-weight distributions between diets was narrow. The interface prominently flagged this limitation, restricted global adjusted ANCOVA outputs to the small overlap window and directed users to causal ANCOVA for a more representative estimand (**Figure 4C**). The conditional comparison at the overlap-based reference body weight yielded adjusted liver means and a weak Diet contrast with wide uncertainty, highlighting that any ANCOVA-style inference here pertains only to a limited subset of animals (**Figure 4C**). Switching to the causal ANCOVA panel, the application estimated total, indirect, and direct effects of Diet using organ-free body weight as mediator by default. Outputs were displayed as posterior means with 95% credible intervals and ROPE probabilities, accompanied by a compact effect plot (**Figure 4C**). In this example, the total loss of liver weight in LPD males was largely mediated by reduced body size, while the direct effect was small and uncertain relative to the ROPE, matching the mediation results reported above (**Figure 4C**). Thus, the application does not silently extrapolate when overlap is poor and provides a defined causal estimand for separating size-mediated from organ-specific components.

Finally, to demonstrate factorial designs, we analyzed a 2 × 2 dataset with Diet (Control vs LowProtein) and Sex (Male vs Female) as grouping factors and Lean mass as outcome. After factor specification, the application constructed the composite Diet × Sex groups and attempted global overlap detection across all four cells. Because no shared trimmed body-weight window existed across the full 2 × 2 design, the application disabled global adjusted means and contrasts and stated that no single reference body weight could be defined without extrapolation (**Figure 4D**). In place of a global conditional comparison, it returned pairwise-matched adjusted means, where each contrast was evaluated at its own overlap-based reference body weight and labelled as such in the plots and summaries (**Figure 4D**). Following the decision framework, the causal ANCOVA module then estimated “Diet” effects separately within each “Sex” level. The application decomposed these stratum-specific effects into total, indirect, and direct components with credible intervals and ROPE probabilities, and the plain-English narrative emphasized whether a meaningful direct organ effect remained after accounting for body size (**Figure 4D**). In this dataset, “Diet” had a strong negative total effect on Lean mass within both Male and Female strata, with sizable negative direct effects in each stratum, indicating Diet-related lean mass loss not fully explained by reduced body weight (**Figure 4D**). This example illustrates the intended behavior in 2 × 2 settings: global conditional effects are refused when unsupported, but stratified causal effects remain available for biologically interpretable inference.

Across these examples, the application reproduces the three analysis branches developed in this work, while enforcing guardrails against ratio-style interpretation and overlap-free extrapolation. It exposes scale choice, overlap diagnostics, reference body weights, and effect decompositions directly in the interface, and returns adjusted effects in original units alongside plain-English summaries to support use by non-specialists.

## 4. Discussion

This work pairs a clear decision framework for organ weight analysis with a Shiny application that makes the framework usable in day-to-day physiology. The application is deliberately narrow in scope: it does not introduce new statistics beyond what we already developed but operationalizes them in a guided workflow. Users load their data, specify the organ, body weight, and one or two grouping factors, and the application then (i) compares RAW ANCOVA to log–log allometry using cross-validated prediction error, (ii) enforces common support via trimmed overlap and a data-derived reference body weight, and (iii) switches to Bayesian causal mediation when size overlap is too weak for a meaningful conditional comparison. The three empirical demonstrations show that the application reproduces the intended estimands: size-adjusted conditional effects when overlap is adequate, and decomposition into total, indirect, and direct effects when body weight is itself altered by treatment.

In our opinion, the main bottleneck is not the absence of appropriate statistical methods, but their limited uptake in routine organ-weight analyses. Ratio normalization remains widespread because it is simple to apply, historically entrenched, and often perceived as an intuitive way to “control” for body size, despite being mathematically unreliable and biologically restrictive in most experimental contexts. These limitations have been documented repeatedly across disciplines for decades, including in physiology and toxicology, yet ratio-based reporting continues to shape standard practice [2–4,9,10]. The purpose of the application is therefore primarily translational: it lowers the practical barrier to principled size-adjusted and causal analyses by providing a guided, user-friendly interface that returns interpretable effects in original units, with explicit uncertainty, and enforces common-support constraints to prevent extrapolation.

Normalization by body weight remains deeply entrenched in physiology, largely because ratio outputs are easy to compute and have long been treated as a default proxy for “relative” organ or metabolic effects. The indirect calorimetry field provides a close parallel: long-standing practice of expressing energy expenditure as kJ per g lean mass or body weight produced biased inference and inflated error, leading to clear recommendations to abandon mass-specific ratios in favor of ANCOVA-style models that condition on body size [14]. Recent methodological discussions in metabolic physiology reiterated that ratio normalization can generate artefactual group differences when treatments shift body mass, and that robust covariate-adjusted models are needed to separate size-mediated from organ-specific effects [15,16]. In that same spirit, CalR was developed as a user-facing tool that makes principled ANCOVA-based analysis routine for calorimetry datasets, lowering the barrier to best practice for non-specialists [17]. Our application extends this logic to organ-weight analysis, where the statistical problem is analogous but where comparable guardrailed tools have been missing.

Indeed, a key contribution of the application is not only automation but guardrails. The overlap check and disabling of global contrasts when no shared support exists are intentional, because extrapolation is exactly how ratio analyses create false certainty. In both the male liver data and the 2×2 lean mass example, the application doesn’t summarize across groups at an unsupported reference size and pushes the user toward an estimand that matches the causal structure of the experiment. This behavior parallels recent efforts to package principled workflows for non-statisticians in adjacent domains, such as toxicology [11]. The broader point is that a friendly interface can still enforce good statistical hygiene if the estimand is made explicit and the software is opinionated about when an analysis is not valid.

There are, of course, limits. First, the application inherits the assumptions of the underlying models. ANCOVA and log–log allometry remain parametric regressions, so strong nonlinearity, heavy tails, or extreme heteroscedasticity may still require specialist handling beyond what the application provides. Second, overlap diagnostics are based on a trimmed rule and cannot replace scientific judgment about whether the remaining common window is biologically representative. The application flags narrow overlap, but it cannot decide what degree of narrowing becomes unacceptable for a given question. Third, mediation decomposes effects only within the assumed causal diagram. If treatment affects unmeasured mediators that also influence organ weight, direct and indirect components should be interpreted as conditional on that simplified structure, not as exhaustive causal truth [5,7,9,11,18]. Thus, the application acts as a guardrail and a facilitator, not a replacement for causal thinking.

Two future directions seem natural. One is extension to other common ratio-like situations already discussed in the manuscript, especially longitudinal pre/post changes where percent-of-baseline shares the same pathologies as organ ratios. A PCA-based standardization module could be added without changing the conceptual logic [5]. Another possible improvement is richer diagnostic guidance that remains simple, so users can spot cases where the workflow is being pushed too far by outliers, non-linear scaling, or batch effects. But any extra features should be added carefully, so the application doesn’t become bloated or give users the impression that everything is automatically trustworthy.

Overall, the Shiny application is a practical answer to a practical problem. Ratios are not “almost fine” or “fine if they look normal”. They are formally unstable, biologically restrictive, and causally opaque. Model-based size adjustment and mediation are not exotic extras; they are the minimum coherent way to ask the questions physiologists actually care about. By embedding these methods in a clean interface with explicit estimands and anti-extrapolation guardrails, we hope to make the statistically sound choice the default choice, and to gradually shift community norms away from per-weight thinking toward explicit size-aware inference.

## CRediT authorship contribution statement

**Lucas Rebiffé:** Conceptualization, Methodology, Software (initial prototype), Validation, Writing – review & editing. **Laurent Gilquin:** Software (refactoring and packaging), Writing – review & editing. **François Leulier:** Writing – review & editing, Supervision, Project administration. **Filipe De Vadder**: Methodology, Software (full implementation and final application), Formal analysis, Visualization, Writing – original draft, Writing – review & editing, Supervision, Project administration.

## Acknowledgements

The authors thank our colleagues for beta-testing OrganScaleR and providing valuable feedback during development.

## Declaration of generative AI and AI-assisted technologies in the manuscript preparation process

During the preparation of this work the authors used Claude (Anthropic) to assist with manuscript editing and figure assembly. After using this tool, the authors reviewed and edited all content as needed and take full responsibility for the content of the published article.

## Data availability

The datasets used in the Results section, the standalone R analysis scripts, and the full OrganScaleR Shiny application (source code and Docker image) are available on Zenodo at https://doi.org/10.5281/zenodo.19188858 (analysis scripts and datasets) and https://doi.org/10.5281/zenodo.20614419 (Shiny application and Docker image). All datasets can be loaded directly into OrganScaleR to reproduce the analyses described in this article.

