## Supplementary material for "OrganScaleR: An Open Shiny Tool for Principled Organ-Weight Inference in Mouse Physiology": Table 1

**Table 1.** Simulation results for organ-to-body-weight ratio, raw ANCOVA, and log–log allometric model across three scenarios (5,000 replicates each, n = 20 per group). True Δ is the between-group difference in organ weight at equal body weight specified by the simulation. Bias is the mean estimation error (estimate minus true Δ). RMSE is the root mean square error of the estimated difference. Rej. rate is the proportion of simulations rejecting the null hypothesis at α = 0.05 (type I error in the null scenario; power otherwise). The allometric exponent was set to 1.6.

|  |  | **Ratio** | | | **Raw ANCOVA** | | | **Log–log allometry** | | |
| --- | --- | --- | --- | --- | --- | --- | --- | --- | --- | --- |
| **Scenario** | **True Δ** (g) | *Bias* (g) | RMSE (g) | Rej. rate | *Bias* (g) | RMSE (g) | Rej. rate | *Bias* (g) | RMSE (g) | Rej. rate |
| Null: body weight shift only | 0 | -0.003 | 0.004 | 0.359 | 0.001 | 0.031 | 0.049 | 0.001 | 0.058 | 0.050 |
| Direct effect only (same body weight) | 0.197 | -0.183 | 0.183 | >0.99 | 0.004 | 0.034 | >0.99 | 0.104 | 0.115 | >0.99 |
| Direct effect + body-weight shift | 0.175 | -0.166 | 0.166 | 0.979 | 0.003 | 0.037 | 0.997 | 0.124 | 0.137 | 0.997 |

*Note: For the ratio method, bias and RMSE are computed after converting ratio-scale estimates to the organ-weight scale for comparability.*
